# Is the “end-of-study guess” a valid measure of sham blinding during transcranial direct current stimulation?

**DOI:** 10.1101/2020.07.11.198416

**Authors:** Christopher Turner, Catherine Jackson, Gemma Learmonth

**Author notes:** These authors contributed equally to this manuscript (Christopher Turner, Catherine Jackson).

## Abstract

Studies using transcranial direct current stimulation (tDCS) typically incorporate a *fade-in, short-stimulation, fade-out* sham (placebo) protocol, which is assumed to be indistinct from a 10-30min active protocol on the scalp. However, many studies report that participants can dissociate active stimulation from sham, even during low-intensity 1mA currents. We recently identified differences in the perception of an active (10min of 1mA) and a sham (20s of 1mA) protocol that lasted for 5 mins after the cessation of sham. In the present study we assessed whether delivery of a higher-intensity 2mA current would exacerbate these differences. Two protocols were delivered to 32 adults in a double-blinded, within-subjects design (*active:* 10min of 2mA, and *sham:* 20s of 2mA), with the anode over the left primary motor cortex and the cathode on the right forehead. Participants were asked “*Is the stimulation on*?” and *“How sure are you?”* at 30s intervals during and after stimulation. The differences between active and sham were more consistent and sustained during 2mA than during 1mA. We then quantified how well participants were able to track the presence and absence of stimulation (i.e. their sensitivity) during the experiment using cross-correlations. Current strength was a good classifier of sensitivity during active tDCS, but exhibited only moderate specificity during sham. The accuracy of the *end-of-study guess* was no better than chance at predicting sensitivity. Our results indicate that the traditional end-of-study guess poorly reflects the sensitivity of participants to stimulation, and may not be a valid method of assessing sham blinding.

## 1. Introduction

Transcranial direct current stimulation (tDCS) is a popular method of neuromodulation that involves the application of weak electric currents to the scalp. These currents are thought to induce temporary changes in the excitability of the underlying cortex, and can subsequently effect changes in behaviours that are regulated by these cortical areas. The majority of clinical and non-clinical studies involving tDCS incorporate some form of placebo control condition into their experimental designs, e.g. the *“fade-in, short-simulation, fade-out”* sham protocol (Nitsche *et al*., 2003; Fonteneau *et al*., 2019). This type of sham protocol is claimed to be indistinguishable from longer, active periods of stimulation (e.g. 10-30 min) on the scalp, and as such, participants are assumed to be blind to the condition that is being administered (Gandiga *et al*., 2006; Poreisz *et al*., 2007; Ambrus *et al*., 2012; Palm *et al*., 2013; Russo *et al*., 2013; Tang *et al*., 2016).

However, a growing number of studies have reported a failure of sham blinding during both high- (2mA) (O’Connell *et al*., 2012; Wallace *et al*., 2016) and lower-intensity (1-1.5mA) tDCS (Goldman *et al*., 2011; Kessler *et al*., 2012; Benwell *et al*., 2015; Greinacher *et al*., 2019; Turi *et al*., 2019). One potential source of these mixed results could stem from the method that is used to assess the success of sham blinding across studies. This measure typically takes the form of a post-stimulation questionnaire to 1) directly probe if participants can identify whether they had received active or sham tDCS (the *“end-of-study-guess”* test) and 2) to quantify the strength of sensations (e.g. tingling or burning) or other side-effects (e.g. changes in mood or concentration levels) experienced during stimulation.

It is assumed that these questionnaires represent a valid measure of sham blinding (Antal *et al*., 2017), yet the responses that are obtained may depend on a range of factors. For example, in the case of multi-session experiments, participants may find it difficult to recall their experiences in prior sessions, particularly when the sessions were many days or weeks apart. They may have taken part in previous electrical stimulation experiments and may know what sensations to expect (Ambrus *et al*., 2012). They may be students who have learned that active protocols tend to be compared to a sham condition, or have gained insight from the experimenter (perhaps disclosure of this information was mandated by an ethics committee) that multiple and different stimulation protocols would be administered. Finally, the questionnaires may not sufficiently clarify technical jargon such as “sham” or “placebo” in lay terms, or may fail to specify whether participants are expected to report only the side-effects that were experienced for a prolonged period of time, or those felt only briefly. In short, the end-of-study questionnaires represent a single snapshot in time - by definition, after the experiment has ended - and yet few studies ask participants to explain the reasons for their choice.

We recently showed in Greinacher *et al*., (2019) that sham blinding was compromised during 1mA tDCS using a novel method of assessment. Participants undertook a simple, forced-choice reaction time task during 10min of 1mA active tDCS and a 20s 1mA sham protocol, based on Minarik *et al*., (2016). None of the participants had received electrical stimulation before, and the two protocols were carefully double-blinded during their application. At regular, 30 second intervals during the 16min experiment participants were asked *“Is the stimulation on?”* (yes/no) and *“How sure are you?”* (0-10 scale). We identified a prolonged period of difference in the perception of active and sham protocols at a group level. Participants were confident that stimulation during the sham protocol switched off after 2mins, whilst they remained confident that stimulation during the active protocol was still switched on until around 11.5mins after onset. We argue that probing the success of sham blinding online, *during* the course of the tDCS delivery, enables researchers to identify differences in perception that might influence the performance of the task that is being undertaken at that point. For example, participants may reduce their effort on the task when they sense the stimulation has ended, leading to differences in performance during the active and sham conditions as a result of failed sham blinding rather than induced neural effects of tDCS.

Our results in Greinacher *et al*., (2019) partially conflict with those observed in Ambrus *et al*., (2012), where multiple (although fewer, at only 7) online probes were presented during stimulation. Participants were asked to report the location and strength of scalp sensations every 1.75min during 10min of 1mA anodal and cathodal tDCS, plus a 30s sham condition. Overall, in line with our results, strength ratings gradually reduced over time, and this reduction was faster in the sham condition compared to anodal and cathodal (the drop was significant by 2.25 min in sham, 4min in anodal and 5.75min in cathodal). However, after excluding the 12 tDCS *investigators* who had participated as subjects, there were no differences in strength ratings between the 3 stimulation protocols in either the naïve or experienced participant groups. These participants were also unable to tell whether they had received real or placebo stimulation. Although our participants were similarly naive to tDCS, we found clear differences in the perception of active anodal and sham stimulation at this low-intensity current strength. However, it should be noted that different questions were used in the two studies which may have contributed to this difference: an analogue scale from *no sensation* to *extreme discomfort* in Ambrus *et al*., (2012), and a binary choice of whether the stimulation was switched on, plus a confidence rating, in Greinacher *et al*., (2019).

In this study we aimed to extend our results in Greinacher *et al*., (2019) by repeating the experiment at a higher-intensity current of 2mA rather than 1mA. We predicted that participants would find it easier to identify the presence and absence of stimulation during both active and sham tDCS when 2mA is delivered, due to stronger sensations on the scalp. Specifically, participants were expected to report that the stimulation was switched on for a longer duration in the active condition compared to our observations during 1mA. Although in Greinacher *et al*., (2019) we found no tDCS-induced changes in reaction time during 1mA stimulation, we hypothesised that a higher, 2mA current may lead to improvements in this outcome measure.

## 2. Method

### 2.1. Pre-registration

The pre-registered study protocol and full datasets are available at https://osf.io/4ath9/.

### 2.2. Participants

Thirty two participants were tested (mean age = 25.69 years, range = 20-41, 22 females). All participants were right-handed with normal or corrected-to-normal vision, had no contraindications to tDCS (Rossi *et al*., 2009), and had not taken part in an electrical stimulation study before. The sample size was based on a planned one-tailed, repeated measures t-test on the reaction time data, where an effect size of *d* = 0.45 was expected (as observed in Minarik *et al*., 2016), power = 0.8 and *α* = 0.05. The study was approved by the University of Glasgow College of Science and Engineering ethics committee and performed in accordance with the Declaration of Helsinki. All participants provided written informed consent.

### 2.3. Transcranial direct current stimulation

A direct current was applied using a battery-driven constant current stimulator (NeuroConn GmbH, Germany). The parameters were identical to those in Greinacher *et al*., (2019), although a 2mA current was delivered here rather than 1mA. Two tDCS protocols (*active* and *sham*) were applied in a double-blinded, counterbalanced, within-subjects design. Double blinding was achieved using the study mode of the NeuroConn stimulator. Pre-allocated 5-digit codes were entered into the device, which initiated either the active or sham protocol. The active protocol involved 10min of 2mA stimulation and sham 20s of 2mA stimulation, with both protocols including an additional 30s ramp-up and 30s ramp-down period. In both protocols, the anode was centred over the left primary motor cortex (C3) and the cathode placed horizontally over the right forehead (Figure 1). Both carbon rubber electrodes measured 5×7 cm, were placed inside 0.9% NaCl saline soaked sponges and were attached to the scalp using rubber bands. The mean impedance at the start of stimulation was 4.27 kΩ (range = 2.2-9.6 kΩ).

**Figure 1.**
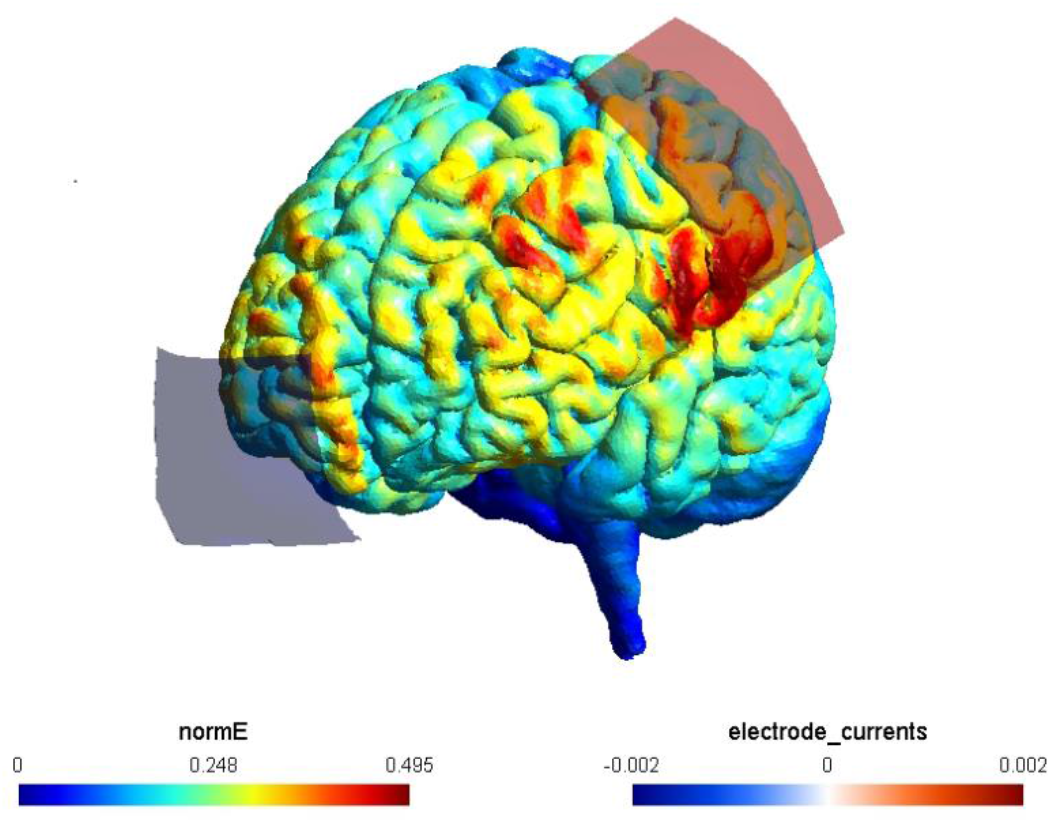
The electrode montage and simulated current flow (SimNIBS 3.1.0, Thielscher et al., 2015). A 5×7cm anode was centred vertically on the left primary motor cortex (C3) and a 5×7 cm cathode was positioned horizontally over the right forehead. The normalised induced electric field (normE) is shown in V/m and the current induced by each electrode in mA.

### 2.4. Behavioural task

Based on Minarik *et al*., (2016), and identical to Greinacher *et al*., (2019), participants completed a simple, forced-choice reaction time task before, during and after stimulation. Stimulus materials are available at https://osf.io/2zwhg/. In Block 1 (baseline) there were 100 trials (lasting around 3.5 minutes), where either a diamond or square appeared in the centre of the screen for 100ms, followed by a fixation cross of variable duration (1700-21000ms). Participants were instructed to respond as fast as possible using their right hand, by clicking the left mouse button to respond to a diamond and the right button for a square. In Blocks 2-4 two probe questions were asked after every 30 s, with each question being presented for a fixed duration of 4500ms. For the first question (*“Is the stimulation on?”*) participants were instructed to click within either the *yes* or *no* box on the screen. For the second question (*“How sure are you?’’*) participants were instructed to click along a visual analogue scale ranging from 0-10, where 0 = very unsure and 10 = very sure. There were 32 probe points in total during Blocks 2-4 spanning a total of 16 minutes.

### 2.5. Procedure

Thirty practice trials were completed, followed by the set of Block 1 trials. The electrodes were then positioned and fixed on the scalp. The resistance was checked and lowered if found to be >10 kΩ by adjusting the rubber bands. At the start of Block 2, the 30s tDCS ramp up was initiated at exactly the same time as the behavioural task was started. Blocks 2-4 were then completed continuously, without breaks, and the end of Block 3 coincided with the end of the 30s ramp-down period in the active condition. Block 4 was completed entirely post-stimulation in both protocols. For each participant, the two sessions were performed at least 24 hours apart. At the end of each session, participants completed a questionnaire which probed their ratings of headache, tingling, itching, burning and pain during stimulation on a scale of 1 = not at all to 5 = very strongly. Finally, they were also asked to guess which of the two sessions had involved sham at the end of the experiment.

## 3. Results

### 3.1. Reaction times

The median reaction time for correct trials was calculated separately for blocks 1-4. The baseline (Block 1) median RT was then subtracted from each subsequent block to create ΔRT values for blocks 2-4. As per the pre-registered analysis plan, a series of 3 one-tailed repeated measures t-tests were performed to compare the ΔRT between the active and sham stimulation conditions in blocks 2, 3 and 4. There were no ΔRT differences between active and sham tDCS in any of the 3 blocks (Block 2: *t*(31) = −0.04, *p* = 0.49; Block 3: *t*(31) = 0.46, *p* = 0.33; Block 4: *t*(31) = −0.18, *p* = 0.43). No differences in ΔRT were observed when comparing the present 2mA dataset and the 1mA data from Greinacher *et al*., (2019) in either the active or sham protocol (2x two-tailed *t*-tests, minimum *p*-value = 0.21).

### 3.2. Effectiveness of sham-blinding during 2mA tDCS

A weighted score was created for each probe point, per participant, where a “yes” response to the question *“Is the stimulation on?”* was assigned a value of +1, and a “no” response a value of −1. This was multiplied by the confidence rating provided in response to the question *“How sure are you?”*, and the weighted scores therefore ranged from +10 = high confidence that the stimulation is on, to −10 = high confidence that it is off. 95% confidence intervals were then bootstrapped for each probe point using 5000 permutations of the data, separately for active and sham (Figure 2).

**Figure 2.**
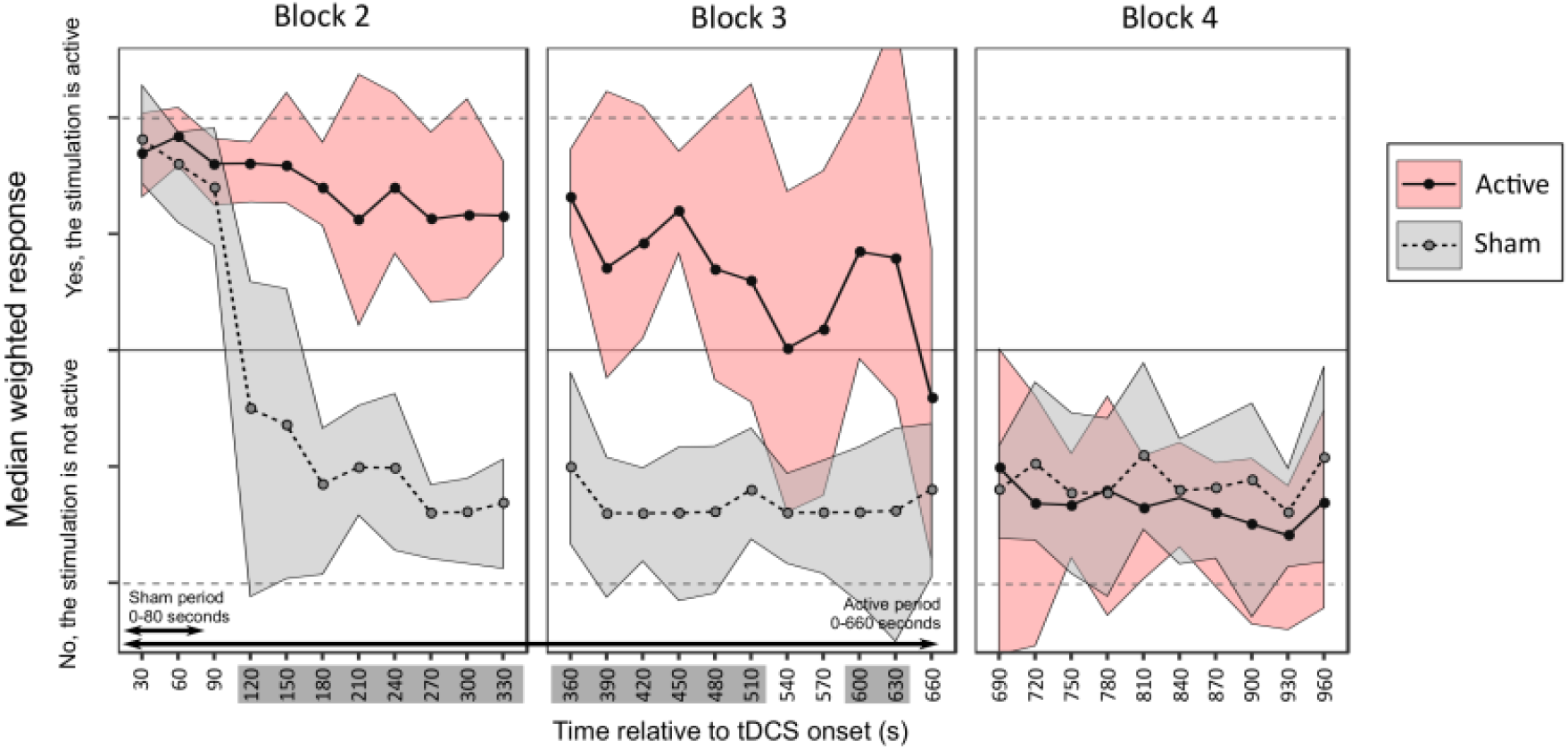
Median weighted scores with 95% confidence intervals for the active (pink) and sham (grey) protocols. The time points with non-overlapping confidence intervals between the two protocols are highlighted in dark grey on the x-axis.

The confidence intervals for active and sham overlapped for the first 90s, where participants reported with high confidence that the stimulation was switched on in both protocols. By 2 mins after onset, participants were confident that the sham stimulation had ended, whereas they reported maintained confidence that the active tDCS was still switched on until 510s (8.5min) after onset and again from 600-630s (10-10.5min) post-onset. By the end of Block 3, at 660s (11min after onset), coinciding with the point at which the active tDCS had fully ramped down, participants reported that the stimulation had ended in both conditions. This was an increase of 3 non-overlapping time points compared with the 1mA dataset in Greinacher *et al*., (2019), although the confidence intervals for the two studies overlapped during both active and sham conditions, mainly due to inter-subject variability.

Wilcoxon signed-rank tests were performed to compare the strength ratings of the 5 sensory side-effects across the two tDCS protocols. Similar to Greinacher *et al*., (2019), itching was stronger in the active condition relative to sham (*Z* = −3.13, *p* = 0.002) but there were no differences for headache, tingling, burning or pain (minimum *p* = 0.197).

### 3.3. Cross-correlation sensitivity analysis

Although the results highlighted clear group-level differences in the time-course of perceptions associated with active and sham tDCS, we noted a high degree of inter-individual variability in the sensitivity of participants to the presence or absence of stimulation. As an exploratory follow-up, we formally assessed this variability using cross-correlations, which can be used to quantify the similarity between two time-series datasets in the form of a correlation coefficient. Specifically, we cross-correlated each participant’s response curve, separately for active and sham stimulation, with the “ideal response” curve for that condition. The ideal response represents the curve that we would expect to observe if the participant was able to report the presence and absence of stimulation with 100% accuracy (see examples of this in Figure 3, right panels). To test the specificity of this analysis method we also performed the cross-correlations with the ideal response for the *opposite* condition (i.e. the response curves for the *active* condition compared to the ideal *sham*, and vice versa). This resulted in a *congruent* and *incongruent* cross-correlation coefficient for each participant during active and sham stimulation, where *r* = 1 represents a perfect positive correlation between the participant’s response and the ideal response.

**Figure 3.**
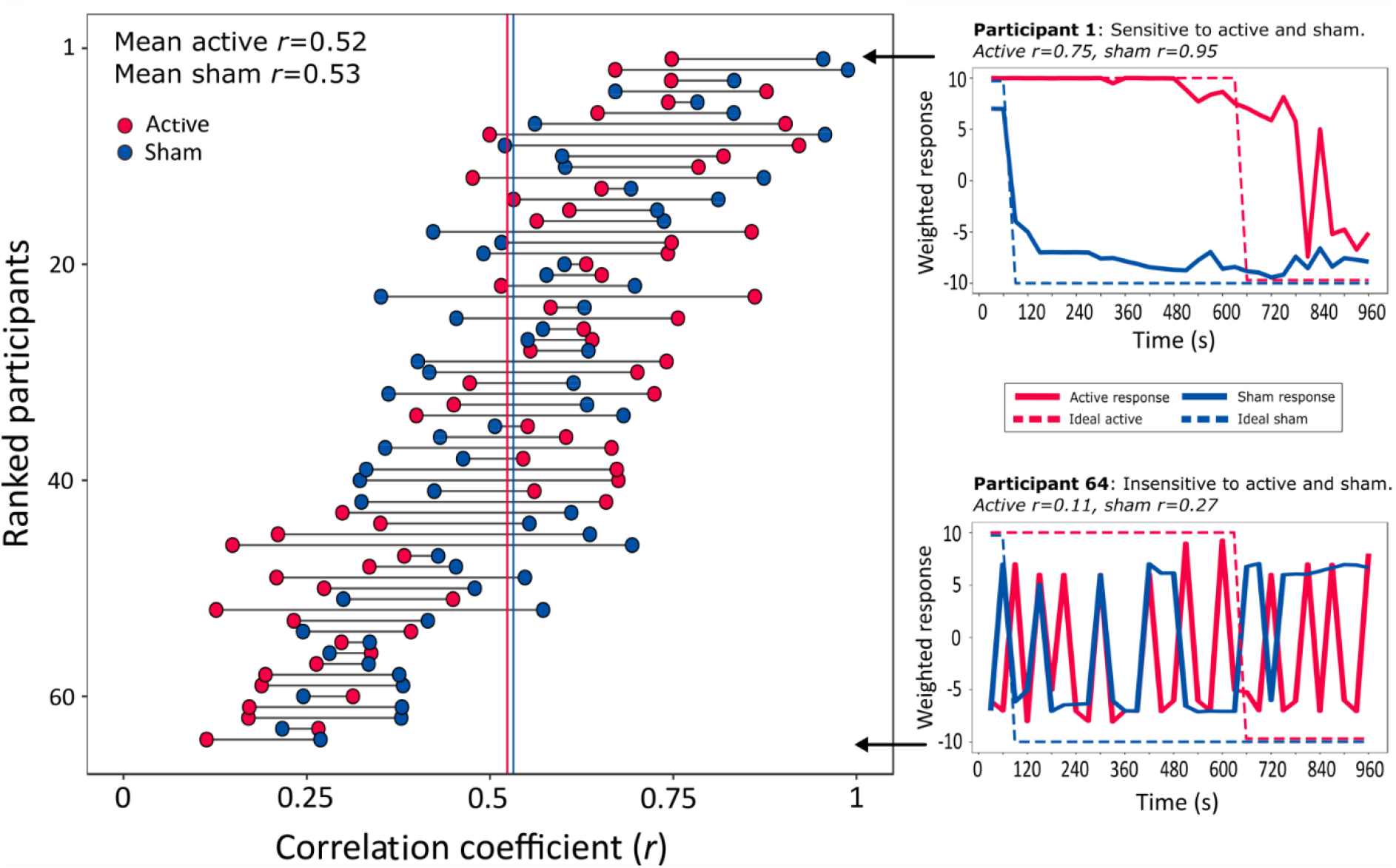
Peak cross-correlation coefficients. The left panel shows all 64 participants ranked by the sum of their Pearson’s correlation coefficients for the active and sham protocols. Mean *r*-values for active and sham are shown as red and blue vertical lines respectively. The panels on the right show the response curves from 2 participants to illustrate the range of sensitivities that were observed across individuals. The highest ranked participant (top right) was highly sensitive to the onset and offset of stimulation during both active and sham protocols. The participant with the lowest sum of coefficients (lower right) responded randomly during both protocols and was thus considered insensitive to the presence of active and sham.

The cross-correlations were performed by first detrending the data vectors then using the *xcorr* function in Matlab with the *‘scaleopt’* parameter set to *‘coeff’*. A sliding window was used because we anticipated that there may be a short delay between the stimulation switching on/off and the participants reporting this change. The maximum Pearson’s correlation coefficient was extracted and used in subsequent analyses. The cross-correlation analysis was performed on both the present dataset obtained during 2mA tDCS and the 1mA dataset from Greinacher *et al*., (2019).

Figure 3 (left panel) shows the peak correlation coefficients of each of the 64 individual participants tested. The mean peak coefficient was moderately large for both the active (*r* = 0.52, *SD* = 0.22) and sham conditions (*r* = 0.53, *SD* = 0.19), with no difference observed between tDCS protocols (paired samples t-test, active *vs* sham: *t*(63) = 0.27, *p* = 0.79). However, there was high variability across participants. The panels on the right of Figure 3 show the data from 2 individual participants: one with high sensitivity to both active and sham (top right panel; *r_active_*= 0.75, *r_sham_*= 0.95), and another individual with low sensitivity to both stimulation conditions (lower right panel; *r_active_*= 0.11, *r_sham_*= 0.27). There was a small within-participant correlation of sensitivity during the active and sham stimulation conditions *r*(64) = 0.26, p = 0.38.

#### 3.3.1. Higher sensitivity to 2mA than 1mA tDCS

The effect of current strength on sensitivity was then assessed (Figure 4, panels A & B). We predicted that sensitivity (i.e. the correlation between the participants’ response and the congruent ideal response) would be higher during 2mA compared to 1mA stimulation due to stronger sensations on the scalp. As expected, a mixed ANOVA (2 x current strengths and 2 x stimulation types) identified a large main effect of current strength *F*(1,62) = 10.62, *p* = 0.002, η_p_^2^ = 0.146. There was no effect of stimulation type *F*(1,62) = 0.07, p = 0.79, η_p_^2^ = 0.001 and no interaction between current strength and stimulation type *F*(1,62) = 1.95, *p* = 0.17, η_p_^2^ = 0.03.

**Figure 4.**
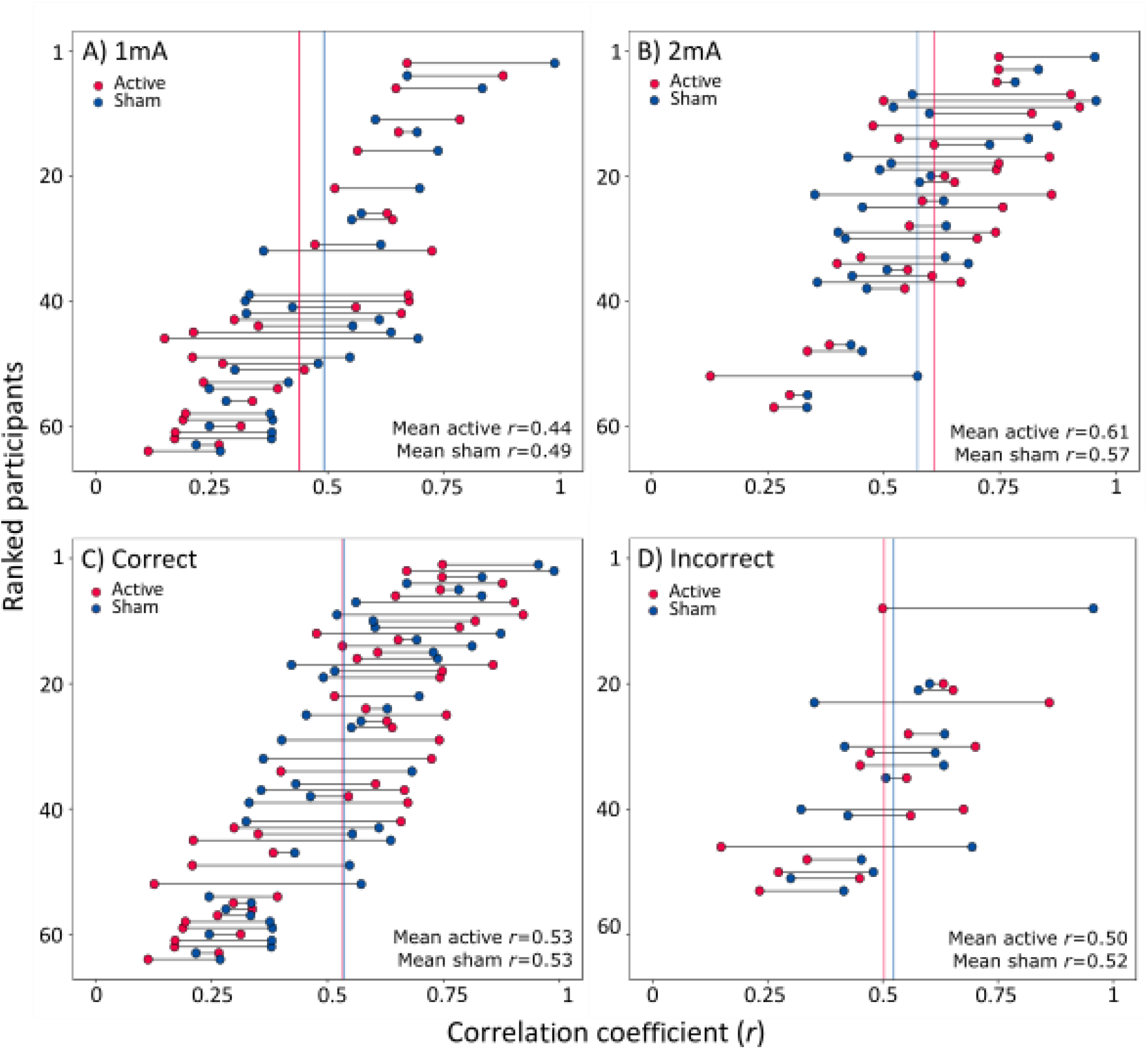
Panels A and B: Peak cross correlations separated by current strength (A = 1ma, B = 2mA). Panels C and D: Peak cross correlations separated by the accuracy of the end-of-study guess regarding which of the two sessions had involved sham (C = correct participants, D = incorrect participants).

#### 3.3.2. No relationship between sensitivity and end-of study guess

We reasoned that this cross-correlation measure of sensitivity should also correspond with the binary response provided at the end of the study, in which participants were asked to guess which of the two sessions had involved sham stimulation (Figure 4, panels C & D). Specifically, participants who correctly identified the sham session were predicted to be able to track the presence and absence of stimulation more closely in both tDCS protocols than those who were incorrect. In total, 75% (48/64) of our participants responded correctly, with similar results during 1mA (25/32 = 78.1%) and 2mA tDCS (23/32 = 71.9%). Of note, there was no overall difference in the sensitivity of participants who guessed correctly compared to those who were incorrect during active stimulation: Welch’s t-test *t*(32.1) = 0.47, *p* = 0.64 or during sham: *t*(29.9) = 0.22, *p* = 0.83. The individuals who guessed correctly had a wide range of sensitivities, some with very high correlation coefficients (max *r* = 0.99) and others with low coefficients (min *r* = 0.11, Figure 4C). Although none of the 16 individuals who were *incorrect* in their end-of-study guess were consistently highly sensitive to both protocols, they did not cluster around the lower correlation coefficients like we had expected (max *r* = 0.96, min *r* = 0.15, Figure 4D).

#### 3.3.3. Specificity analysis

In addition to indexing the sensitivity of participants to the presence of stimulation, we also assessed the specificity of this analysis method. A 2*x*2 ANOVA (2 stimulation types and 2 congruence types) identified a main effect of correlation congruence (*F*(1,63) = 55.87, *p* < 0.001, η_p_^2^ = 0.47), indicating that response curves were specific to the stimulation protocol that the participant was receiving (mean *congruent: r* = 0.53, 95% *CI* = [0.49, 0.57] and *incongruent: r* = 0.39, 95% *CI* = [0.37, 0.41]). There was no main effect of stimulation type (*F*(1,63) = 2.14, *p* = 0.15, η_p_^2^ = 0.33) and no stimulation *x* congruence interaction (*F*(1,63) = 0.28, *p* = 0.6, η_p_^2^ = 0.004).

Finally, receiver operating characteristic (ROC) curves (Figure 5) were used to assess the specificity of the 3 binary-outcome classifiers described above: a classification of the correlation coefficients as belonging to 1) the congruent or incongruent correlation analysis, 2) the 1mA or 2mA dataset, or 3) the participants who were correct or incorrect in their end-of-study guess. The area under the curve (AUC) provides a measure of how well a model can distinguish between two states, where a value of 1.0 represents a perfect classifier and 0.5 represents random chance.

**Figure 5.**
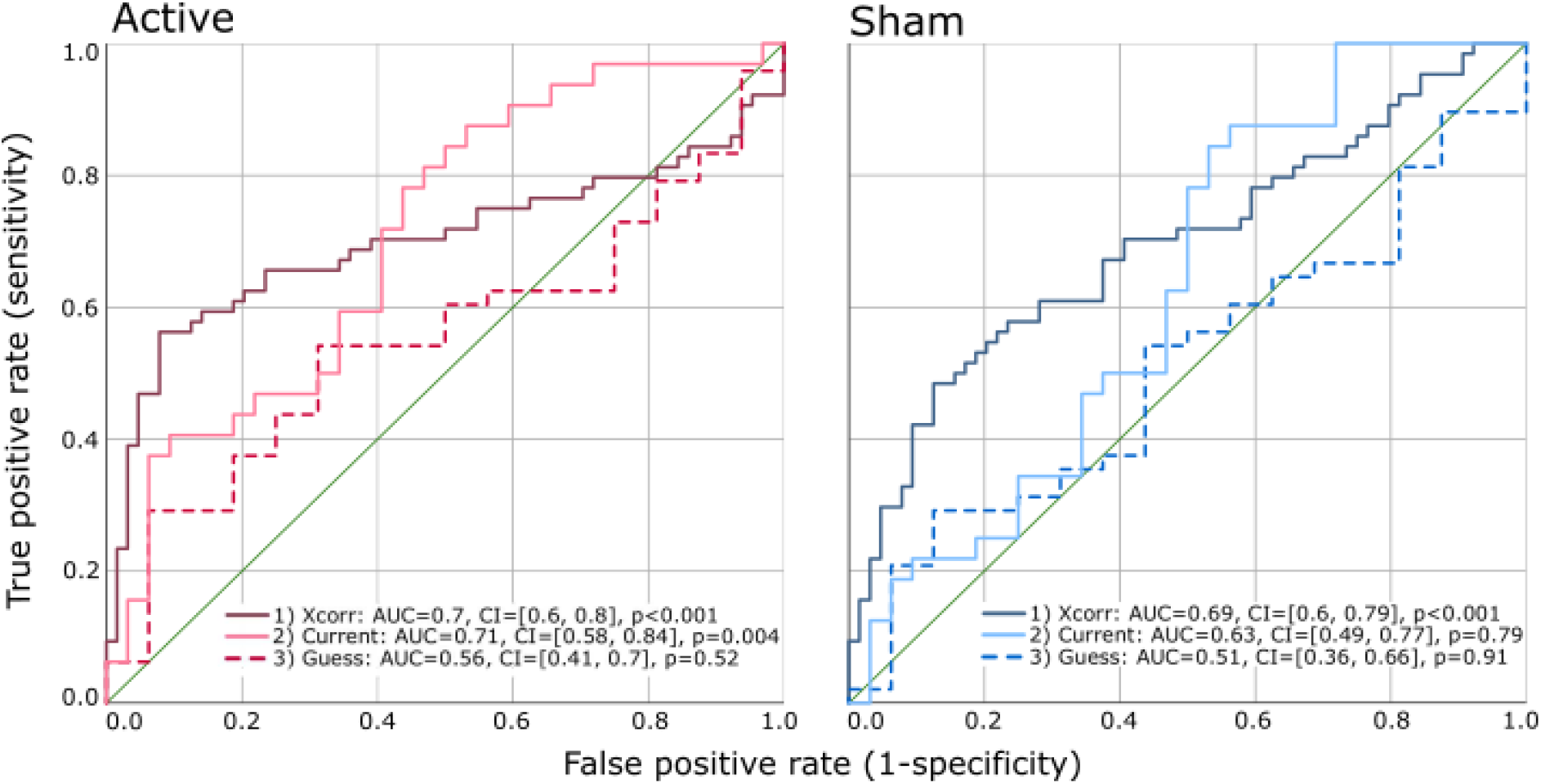
ROC curves demonstrating the sensitivity and specificity of 3 classification methods for the active and sham tDCS protocols: 1) cross-correlations, 2) current strength and 3) the end-of-study guess.

##### 1) Cross-correlation congruence

The ROC curve indicates that congruence is a good predictor of correlation strength (sensitivity), with a classification accuracy of around 70% (active: AUC = 0.7, 95% *CI* = [0.6, 0.8], *p* < 0.001; sham: AUC = 0.69, 95% *CI* = [0.6, 0.79], *p* < 0.001).

##### 2) Current strength

Current strength is a good predictor of sensitivity, but only during active stimulation, with an accuracy of 71% (AUC = 0.71, 95% *CI* = [0.58, 0.84], p = 0.004). Current strength is a less effective predictor of sensitivity during sham stimulation (AUC = 0.63, 95% CI = [0.49, 0.77], *p* = 0.079).

##### 3) End-of-study guess

The accuracy of the end-of-study guess failed to predict sensitivity in either stimulation condition, with only random-chance levels of accuracy (active: AUC = 0.56, 95% *CI* = [0.41, 0.7], *p* = 0.52; sham: AUC = 0.51, 95% *CI* = [0.36, 0.66], *p* = 0.91).

## 4. Discussion

We have demonstrated that participants are able to identify that a 10min 2mA tDCS current is being delivered for a longer period of time compared to a 20s 2mA sham protocol. As predicted, the perceived differences between active and sham tDCS were more pronounced during high-intensity 2mA stimulation compared to the 1mA current applied in Greinacher *et al*., (2019), with an additional 3 time points with non-overlapping confidence intervals in the second half of the 10min stimulation period during 2mA tDCS. Also aligned with our previous findings, we found no beneficial effects of 10min of anodal stimulation over the left primary motor cortex in improving reaction times. Using a novel method of quantifying the sensitivity of participants using cross-correlations, we found that current strength was a good classifier of sensitivity during active tDCS, but was only a moderately accurate classifier during sham. Finally, the accuracy of the binary end-of-study guess was no better than chance at classifying sensitivity during either active or sham stimulation, raising important questions regarding the validity of this method of assessing sham-blinding.

We were able to uniquely quantify the sensitivity of participants to the presence of active and sham stimulation throughout the course of the experiment by using cross-correlations. Participants with large correlation coefficients were considered to be sensitive in their ability to track the presence (and absence) of stimulation, and those with small coefficients as insensitive. This method adds key information regarding the time-course of sham blinding which is unavailable to researchers using only the standard methods of assessing placebo control, namely administering questionnaires at the end of the study. These post-study questionnaires usually incorporate a binary probe question to assess whether participants can identify if an active or sham protocol was delivered, or which of multiple sessions had involved active or sham, in the case of repeated measures designs. Since our cross-correlation measure quantified how well participants could track the presence of stimulation during both 10min of active anodal and a 20s sham, we expected to observe an association between this measure and the accuracy of their end-of study guess. Specifically, that a high sensitivity would lead to an increased likelihood of correctly dissociating the two conditions. On the contrary, our results showed no association between the end-of-study guess accuracy and sensitivity to either active or sham tDCS, and the end-of-study guess was, in fact, a poor classifier of sensitivity.

The end-of-study questionnaire method of assessing sham blinding is undoubtedly the most popular method used in electrical brain stimulation studies (Antal *et al*., 2017). However, if as we show here, the end-of-study guess does not reflect the participants’ sensitivity to the presence of stimulation, what might this guess represent? There are many potential explanations for this discrepancy, such as the presence of confusing and technical terminology in the questionnaires (e.g. did participants understand the distinction between “sham” and “active”, and did they know whether to report fluctuating and/or sustained scalp sensations?, perhaps a poor memory for prior sessions, and the influence of prior experience or knowledge of tDCS methodology, or even subtle priming by the experimenters (Rabipour *et al*., 2018, 2019). All of these variables could feasibly influence the outcome of a reflective binary probe question and these factors should now be examined systematically in further studies.

It is important to note that although the congruence of the cross-correlations was a better classifier of participants’ sensitivity to stimulation than the end-of-study guess, it was only accurate in around 70% of individuals. This serves to highlight a moderate degree of inter-individual variability in sensitivity across our 64 participants. Although the mean coefficient for congruent correlations was *r* = 0.53, this ranged from as low as *r* = 0.11 to as high as *r* = 0.99. We also found that sensitivity was relatively consistent within each individual across the active and sham protocols (Fig 3, left panel), with a small correlation of *r*(64) = 0.26, *p* = 0.38. For example, participants who had a high (or low) sensitivity to active stimulation also tended to have a high (or low) sensitivity to sham. This points to the presence of sub-groups of individuals, where some people are more, or less, susceptible to sham-blinding than others. It may therefore be possible to screen individuals and select those with low sensitivity during the recruitment process for research studies in order to maximise the likelihood of blinding the conditions successfully. However, this is unlikely to be sustainable when recruiting participants for clinical trials, particularly for conditions with a small patient pool, as this would result in the exclusion of a large proportion of otherwise-eligible individuals.

It is perhaps unsurprising that we identified a greater sensitivity to high-intensity (2mA) tDCS compared to low-intensity (1mA) stimulation. At the group-level, participants reported a clear sustained difference between active and sham that was maintained until almost the end of the 11min total stimulation period. In Greinacher *et al*., (2019), during 1mA tDCS, this divergence lasted until 6mins post-onset of stimulation, with sporadic differences lasting until 11.5mins. In fact, the current strength in the present study was a good classifier of sensitivity during the active stimulation protocol (where sensitivity was higher during 2mA than 1mA), but was less successful at predicting sensitivity during sham. In other words, it may be that the scalp is generally most sensitive during the first 1-2 minutes after stimulation onset (coinciding with the sham stimulation period), and this sensitivity is relatively independent of the strength of the current that is applied. In the active condition, the higher-intensity current applied for a longer period results in a prolongation of the induced sensations compared to low-intensity stimulation, thus resulting in participants being able to track the presence of stimulation more effectively throughout the duration of the experiment.

We must acknowledge that our results, at present, only pertain to the specific experimental design, parameters and participant group tested here, and must now be replicated with different electrode sizes, montages, with older people and patient groups, and in experiments adopting a between-group design. It could be that between-group designs result in less pronounced differences between protocols, particularly if participants are naive to tDCS, however, repeated measures designs are commonly used in tDCS research, and are indeed preferential because of their increased statistical power. This is particularly important in a field where observed effect sizes are typically small. Secondly, as noted in Greinacher *et al*., (2019) we probed sham-blinding regularly throughout the study and in doing so we may have drawn attention to the sensations more than in studies using only the traditional end-of-study guess questionnaire. However, in this scenario we would have expected to observe more consistently higher sensitivities across participants than we did, rather than the range (and potential subgroups) of sensitive and insensitive individuals. As we noted in Greinacher *et al*., (2019) we also found that reaction times during 1mA tDCS were no different to those obtained in Minarik et al., (2016) where no online probes were used, indicating that participants were not unduly distracted from the reaction time task by the presence of the online probe questions.

Finally, these results prompt us to make a number of recommendations to improve sham blinding in future studies. Firstly, we recommend introducing at least some online probe questions during the course of stimulation to assess the effectiveness of the placebo control that is being delivered. We also recommend eliminating the traditional *fade-in, short stimulation, fade-out* sham protocol where possible, and adopting “active controls” instead. For example, delivering active stimulation over a cortical site that is thought not to be involved in the behaviour of interest, or alternatively by applying an anodal compared to a cathodal protocol. In some cases this approach may not be viable, for instance in patients after stroke, where it may prove detrimental to induce a further reduction (or indeed an increase) of neural activity. Topical anaesthetics should also be used where possible to minimise or eliminate sensory side-effects (McFadden *et al*., 2011; Guleyupoglu *et al*., 2014). Recently, Neri *et al*., (2020) developed a novel method of sham blinding (*ActiSham*) which involves deliberately shunting the current across the scalp in a controlled manner using multifocal tDCS. This method is claimed to provide similar scalp sensations to active tDCS, without inducing a neural effect, and participants are reportedly unable to dissociate *ActiSham* from active stimulation. It would be beneficial to develop such protocols using the standard, large electrode montages which are used more widely in tDCS research. Further studies should also be carried out to quantify the time course of more specific side effects, such as headache and tingling, and whether they might predict sensitivity to stimulation in a similar manner (Fertonani *et al*., 2015).

## 5. Conclusion

High-intensity 2mA tDCS current results in a more prolonged and consistent period of time where the active stimulation is perceived as being switched on, compared to low-intensity 1mA stimulation. The accuracy of the end-of-study guess (i.e. the standard post-study questionnaire method of determining whether participants can dissociate the active from the sham session) was no better than chance at predicting the sensitivity of participants to the presence of either anodal or sham tDCS. This raises important questions regarding the validity of the questionnaire-based method of assessing sham-blinding.

